# Influence of the artificial sodium saccharin sweetener Sucram^®^ on the microbial community composition in the rumen content and attached to the rumen epithelium in dairy cattle: A pilot study

**DOI:** 10.1101/2020.05.22.110429

**Authors:** Lucas R. Koester, Chiron J. Anderson, Bienvenido W. Cortes, Mark Lyte, Stephan Schmitz-Esser

**Author notes:** **Corresponding author:** Dr. Stephan Schmitz-Esser, Iowa State University, Department of Animal Science, Ames, IA, USA.

## Abstract

The products of rumen microbial fermentations are considered essential for animal growth and performance. Changes in these microbial communities can have major effects on animal growth and performance. Saccharin-based artificial sweeteners can be included in livestock diets to increase palatability and encourage feed intake. Despite the importance of the rumen microbial fermentation, little or no research is available regarding how saccharin-based artificial sweeteners affect rumen content and rumen epithelial microbial communities. The aim of this study was to identify changes in both the rumen content and rumen epithelial microbial communities in response to the supplementation of Sucram^®^, a sodium-saccharin-based sweetener (Pancosma S.A./ADM Groups, Rolle, Switzerland) during standard, non-stress conditions using 16SrRNA gene amplicon sequencing.

The rumen epithelial and rumen content microbiota of five Holstein-Friesian milking dairy cattle were compared before (baseline, BL) and after a 28-day supplementation of Sucram^®^. Illumina MiSeq-based 16S rRNA gene sequencing was conducted, and community analysis revealed significant changes in the abundance of specific phylotypes when comparing BL to Sucram^®^ experimental groups. Sucram^®^ did not have a significant effect on overall rumen microbial community structure between experimental groups. Statistically significant changes in microbial community composition following Sucram^®^ supplementation were observed most consistently across a number of bacterial taxa in the rumen epithelium, while fewer changes were seen in the rumen content. Predicted genomic potentials of several significantly different OTUs were mined for genes related to feed efficiency and saccharin degradation. Operational taxonomic units (OTUs) classified as *Prevotella* and *Sharpea* were significantly (p<0.05) increased in samples supplemented with Sucram^®^, whereas a reduction in abundance was seen for OTUs classified as *Treponema*, *Leptospiraceae*, *Ruminococcus* and methanogenic archaea. This is the first study to report an effect of Sucram^®^ on ruminant microbial communities, suggesting possible beneficial impacts of Sucram^®^ on animal health and performance that may extend beyond increasing feed palatability.

## Introduction

Sucram^®^ (Pancosma, S.A./ADM Groups, Rolle, Switzerland), is a sodium-saccharin-based sweetener which has been approved for use in a range of livestock species including milking dairy cattle. A number of recent publications have described the effect of Sucram^®^ on growth performance in cattle [1, 2], and suggest increases in average daily gain and feed intake during stress conditions when feed is supplemented with Sucram^®^. The effects of artificial sweeteners such as Sucram^®^ on mammalian GI microbiomes have increasingly been studied over the last few years as a possible mechanistic pathway by which these food additives may influence host physiology. Several studies have described alterations in the microbiome caused by the inclusion of artificial sweeteners in monogastric animals and humans [3–11]. However, little to no research into if saccharin-based artificial sweeteners affect the GI microbiota of ruminants is currently available.

The microbial communities in the gastrointestinal tract (GI) of mammals have profound effects on health and performance. GI tract microbial communities of ruminants perform important roles in host metabolism (i.e. cellulose degradation) [12] and are largely distinct from those found in monogastric species. Rumen microbial fermentation provides key metabolic products for the host animal such as short chain fatty acids (SCFA) and vitamins via the breakdown of cellulose, hemicellulose, pectin and other ingested feed [13, 14]. These microbial metabolic products are absorbed directly by the host through the rumen epithelial tissue. More generally, the integrity of the rumen epithelial tissue is essential for animal health and performance as a decrease in rumen epithelial tissue integrity can result in inflammation and symptoms referred to as “leaky gut” [15, 16]. Modulation of the GI tract microbiome to improve ruminant livestock feed efficiency and health is of great interest to livestock producers, and as antibiotics usage in the livestock industry decreases, alternative feed additives, such as pre- and probiotics, are being explored. Artificial sweeteners have not been examined in this context until now.

Rumen microorganisms have been characterized as two main groups; rumen epithelial microorganisms and rumen content microorganisms [17–21]. Currently, there are very few studies that focus on the rumen epithelium microbial communities and their potential effects on the host. Rumen epithelial microorganisms are often considered less transient than their luminal counterparts, suggesting a more stable community that possibly interacts with the host through a variety of yet undiscovered ways [20]. As the rumen epithelium is a major site of nutrient exchange between the rumen content and the animal host, the close proximity of the rumen epithelium microbial communities to the host tissue suggests the rumen wall microbes may influence nutrient exchange or signaling to the host [21, 22]. Additionally, the rumen epithelial microbial communities may play an additional role in important metabolic functions such as nitrogen metabolism, sulfate reduction, and oxygen scavenging [23]. Rumen content microbial communities include the microorganisms attached to particulate matter and those who are planktonic within the liquid phase of the rumen content. These microorganisms are known to be integral for fiber-degradation and feed digestion [24, 25]. Each of these distinct groups are key to metabolic processes in the host, and the differences between the two should be considered when conducting a rumen microbial analysis.

Until now, the effect of Sucram^®^, or that of any other sweetener-based food additive, on rumen microbiota composition has not been analyzed. This study aimed to provide preliminary data to determine changes in the rumen content and rumen epithelium bacterial communities, in response to supplementation of Sucram^®^ in dairy cows under standard, non-stressful, physiological conditions. As a pilot study, targeting possible effects of Sucram^®^ on microbial community composition, we did not aim for identifying possible effects of Sucram^®^ on feed intake or feed efficiency. Given that the ruminant microbiome is critical to animal health and performance and that there is very little general knowledge of microbial organisms inhabiting the rumen epithelium, exploring the effects of Sucram^®^ on ruminant microbial communities may lead to improved understanding of the factors which influence feed efficiency and ultimately lead to better animal health.

## Materials and methods

### Ethics statement

All animal procedures in this study were conducted under approval of the Animal Care and Use Committee at Iowa State University (ISU) (IACUC# 1-18-8670-B).

### Animal trial

Five rumen fistulated, lactating Holstein-Friesian dairy cows housed at the Iowa State University (ISU) dairy teaching farm were included in this trial. To study the effect of the artificial sweetener Sucram^®^ C-150 (Pancosma S.A./ADM Groups, Rolle, Switzerland) on rumen content and rumen epithelium microbial communities, each animal was sampled before (baseline, BL) and after 28 days of Sucram^®^ C-150 feeding. In this way, each animal acted as its own control (BL, pre-exposure to Sucram^®^ C-150) when analyzing potential effects of the compound on rumen microbial communities. Animals were housed together under identical conditions at the ISU dairy farm. All cows received the ISU dairy farm regular diet comprised of ground corn, soybeans, cottonseed hulls, corn silage, baleage and alfalfa hay (53.8% dry matter (DM), 9.46% crude protein (CP), and 13.91% neutral detergent fiber (NDF)). Details of the analysis and chemical composition of the diet are given in Supplementary Table S1. The Sucram^®^ experimental group cows were given 2 grams of Sucram^®^ suspended in 10 ml of 1x sterile phosphate buffered saline (PBS) (final concentration: 0.2g/ml) per day and cow as per feeding protocols provided by Pancosma. The Sucram^®^ C-150 containing solution was added directly through the fistula to ensure all cows consistently received the same amount of sweetener. Our main aim was to identify if the presence of Sucram^®^ has an influence or rumen microbial communities, and due to the administration procedure of Sucram^®^ through the fistula, any sensory stimuli (i.e. taste or smell) would be limited in this experimental setup. Consequently, Sucram^®^ would not have an effect on feed intake as it wasn’t mixed with the feed by-passing any sensory stimuli. Thus, we did not measure feed intake, average daily gain or milk yield in response to administration of Sucram^®^ for this study.

Two sample types were taken for this trial: rumen content, and rumen epithelium biopsies, taken from the dorsal part of the rumen wall, both collected directly through the fistula. Rumen epithelium biopsies were collected using Chevalier Jackson forceps from three locations (separated vertically by 10 cm) in the dorsal rumen sac and were combined to provide a more representative sample of rumen wall microbial communities. Rumen epithelial biopsy samples were briefly rinsed in sterile 1x PBS, immediately snap frozen in liquid nitrogen on site and placed in sterile, 2 ml screw-top centrifuge tubes. Approximately 12 ml of rumen content was collected in 15 ml sterile conical tubes and frozen on site on dry ice immediately after retrieval. All samples were stored at −80°C after collection.

Rumen epithelium and content samples were thawed, and genomic DNA was extracted from approximately 0.1 grams of rumen epithelial sample and approx. 0.2 grams of rumen content sample using the Qiagen DNeasy Powerlyzer Powersoil kit following the manufacturer’s instructions. Mechanical cell lysis was performed using a Fischer Scientific Beadmill 24, and DNA concentrations were determined using a Qubit 3 fluorometer (Invitrogen, Carlsbad, CA, USA).

After extraction, DNA concentrations were adjusted to 25 ng/μl and sent to the ISU DNA facility for sequencing using the Illumina MiSeq platform (Illumina, San Diego, CA, USA). Briefly, the genomic DNA from each sample was amplified using Platinum™ Taq DNA Polymerase (Thermo Fisher Scientific, Waltham, MA, USA) with one replicate per sample using universal 16S rRNA gene bacterial primers [515F (5′-GTGYCAGCMGCCGCGGTAA-3′; [26]) and 806R (5′-GGACTACNVGGGTWTCTAAT-3′; [27])] amplifying the variable region V4, as previously described [28]. All samples underwent PCR with an initial denaturation step at 94°C for 3 min, followed by 45 seconds of denaturing at 94°C, 20 seconds of annealing at 50°C, and 90 seconds of extension at 72°C. This was repeated for 35 total PCR cycles and finished with a 10 minute extension at 72°C. All the PCR products were then purified with the QIAquick 96 PCR Purification Kit (Qiagen, Hilden, Germany) according to the manufacturer’s instructions. PCR bar-coded amplicons were mixed at equal molar ratios and used for Illumina MiSeq paired-end sequencing with 150 bp read length and cluster generation with 10% PhiX control DNA on an Illumina MiSeq platform at the ISU DNA facility.

### Sequence analysis

Sequence analysis was done with Mothur V1.40.5 following the Mothur MiSeq Standard Operating Procedure [28]. Barcode sequences, primers and low-quality sequences were trimmed using a minimum average quality score of 35, with a sliding window size of 50 bp. Chimeric sequences were removed with the “Chimera.uchime” command. For alignment and taxonomic classification of operational taxonomic units (OTUs), the SILVA SSU NR reference database (V132) provided by the mothur website was used. Sequences were clustered into OTUs with a cutoff of 99% 16S rRNA gene similarity (=0.01 distance). As stated above, due to the clear difference between rumen content and rumen epithelial microbial communities, rumen content samples were analyzed separately from the rumen epithelial samples.

To compare alpha diversity between experimental groups, reads were randomly subsampled to accommodate the sample with the lowest number of reads across data sets (20,000 sequences for rumen content samples and 20,000 sequences for rumen epithelial samples). Measurements of Chao species richness, Shannon Diversity, and Simpson evenness were taken to compare community structures between experimental groups. The means of the experimental group alpha diversity measures were compared using a pooled t-test assuming equal variance. Because the analysis compared cattle rumen samples of the same type (rumen content/rumen content or rumen epithelial/rumen epithelial) and because animals of the same group at the same farm, the samples were assumed to be highly similar in nature. Therefore, Bray-Curtis was selected as the dissimilarity coefficient because of its ability to compare closely related samples. After dissimilarity coefficients were assigned to each sample, experimental groups were compared using the analysis of similarity (ANOSIM) package provided by mothur.

All plotting was completed using ggplot2, v2_3.1.1 graphing package [29, 30] in R 3.6.0. Overall variation in bacterial communities was visualized using principle coordinate analysis (PCoA). Canonical correlation analysis (CCA) was used to visualize the variation strictly due to Sucram^®^. This information was generated with the Phyloseq (v1.28.0 [31]) and Vegan (v2.5-5, [32]) packages using the shared and taxonomy file generated in mothur. Sequences were randomly subsampled to 20,000 sequences and Bray-Curtis dissimilarity measures were used to generate distances between samples for the PCoA and CCA plots.

Differences in individual OTUs were compared using Linear Discriminant Analysis (LDA) Effect Size (LEfSe, [33]), identifying OTUs that most likely explain the greatest between-group variation. LEfSe performs a Kruskall-Wallis test to analyze all OTUs, broadly selecting OTUs that show the most variation between sample types. The retained features then undergo a pairwise Wilcoxon test, removing any OTUs that do not differ in ranking. In the last step, a linear discriminant analysis model is built from the retained OTUs to determine the effect sizes for each feature. P-values of <0.05 were considered significant.

### Predictions of microbial functional potential based on a rumen genome collection

We combined assembled draft genomes of rumen content organisms from three studies that used both metagenome shotgun sequencing data[34], as well as whole genome sequencing of cultured organisms [35] to create a rumen genome collection (RGC). We then compared the representative 16S rRNA gene sequences from the 100 most abundant OTUs in both the rumen content and rumen epithelium datasets generated in this study to the RGC using BLAST+ [36], with a threshold of 97% sequence similarity. The taxonomic information from the RGC for each match was then appended to the taxonomic information provided by Silva and NCBI Blast. This was done to offer additional, possibly more accurate taxonomic classification for these sequences, as well as provide some information about the possible genetic potential of these OTUs. The genomes from the RGC that matched the 16S rRNA gene sequences of the significant OTUs were then uploaded into PATRIC (v3.6.3 [37]) and annotated. The genetic features of each genome identified in the annotation were then mined for genes involved in feed efficiency and the degradation of saccharin based on the work done by Shabat et al [38] and Deng et al [39]. Although the genes involved in degradation of saccharin are not known, Deng et al predicted genes within the aromatic hydrocarbon degradation and dissimilatory sulfate and sulfide oxidation pathways to be important for saccharin degradation (Supplementary Table S2).

### Data availability

The 16S rRNA gene sequences have been submitted to the NCBI Sequence Read Archive SRA and are available under the BioProject ID PRJNA554894.

## Results

### Rumen content microbial communities

3,291 OTUs were generated from rumen content samples after quality control and removal of OTUs representing less than 10 sequences from the 910,228 high quality sequences from 10 samples. The average sequencing depth per sample was 91,022 sequences with a standard deviation of 32,011 sequences. 99.5% of the reads were bacterial and 0.5% were archaeal. The 3,291 OTUs were assigned to 20 phyla with *Bacteroidetes*, *Proteobacteria* and *Firmicutes* being the most abundant: representing 39%, 30% and 20% of all reads, respectively (Fig. 1). Within the rumen content OTUs, OTU 1, classified as *Ruminobacter* RUG14687 with 100% sequence similarity using the RGC and *Succinivibrionaceae*_UCG-001 (88.98% sequence similarity, Silva v132), was the most abundant and accounted for 25.6% of all reads and had a 100% sequence similarity with OTU 1 of the rumen epithelium dataset. Among the 50 most abundant rumen content OTUs, 27 OTUs were classified within the family *Prevotellaceae*, a family which accounted for 32% of all reads from the rumen content data set. A list of the 50 most abundant rumen content OTUs can be found in Supplementary Table S3. Highly abundant genera within the rumen content include: *Succinivibrionaceae*_UCG-001 (25.6%), *Prevotella*_1 (18.3%), *Treponema_2* (3.3%), *Rikenella* (2.1%) and *Fibrobacter* (1.8%) (Fig. 2).

**Fig. 1.**
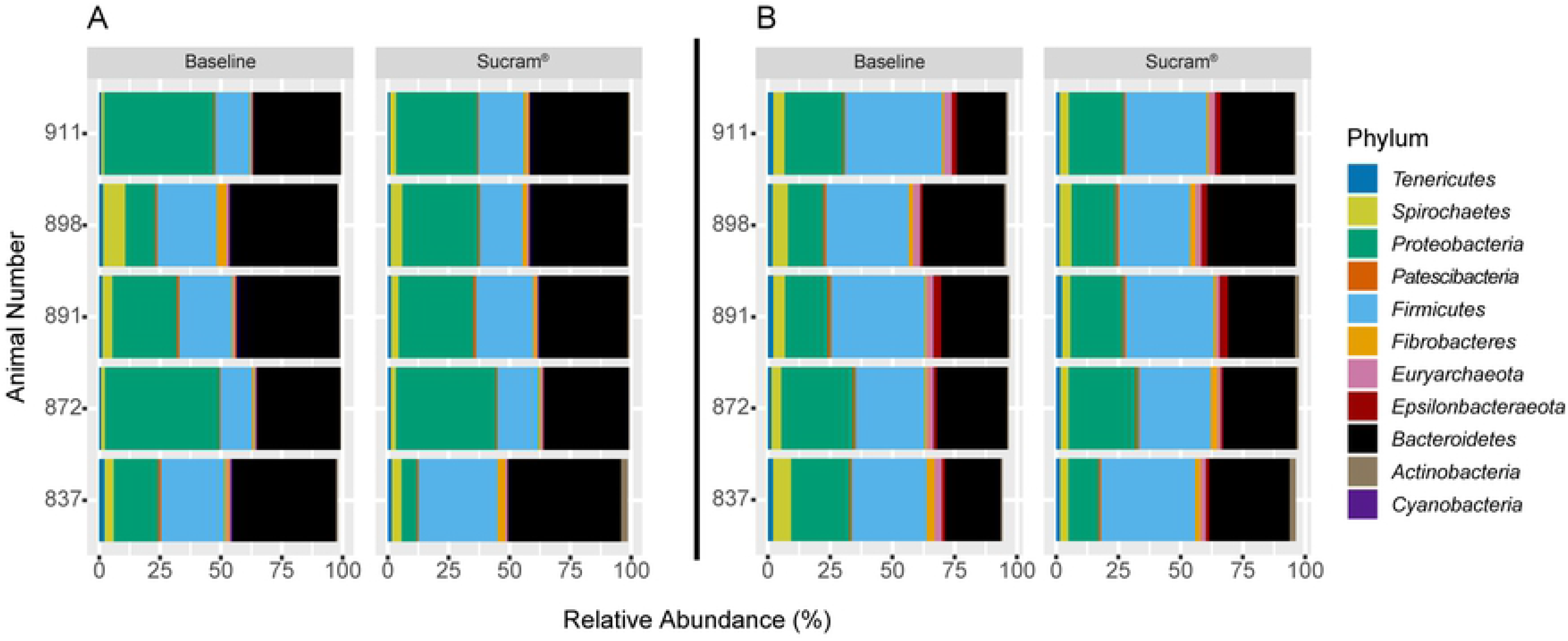
Relative abundance of rumen content (A) and rumen epithelium (B) microbial communities on phylum level in response to Sucram ^®^ addition. Data are shown for baseline (before) and after 28 days of Sucram^®^ addition to the diet. Only the 10 most abundant phyla per experimental group are shown.

**Fig. 2.**
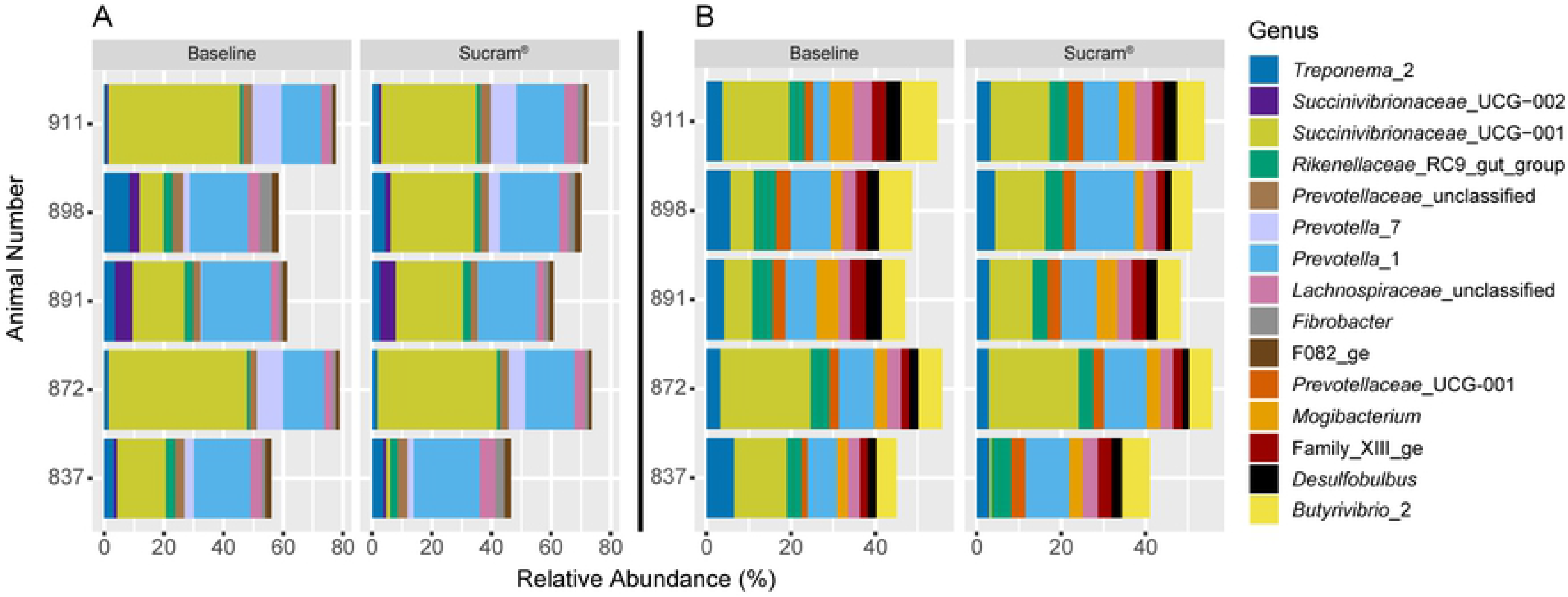
Relative abundance of rumen content (A) and rumen epithelium (B) microbial communities on genus level in response to Sucram ^®^ addition. Data are shown for baseline (before) and after 28 days of Sucram^®^ addition to the diet. Only the 10 most abundant genera per experimental group are shown.

No significant differences were observed in microbial diversity (Shannon, *P* = 0.9), species richness (Chao, *P* = 0.81) and microbial community evenness (Simpson, *P* = 0.73) when comparing BL and Sucram^®^ rumen content experimental groups (Supplementary Table S4). BL and Sucram^®^ bacterial communities of the rumen content were compared using ANOSIM, and no significant differences between experimental groups were found (p = 0.527, R =-0.036). PCoA plots generated with these data also provided no evidence of community clustering according to experimental group and 10.5% of the total variation was due to experimental group (CCA, Fig. 3). The lack of clustering and low amount of variation due to experimental group corroborates the reported ANOSIM community comparison.

**Fig. 3.**
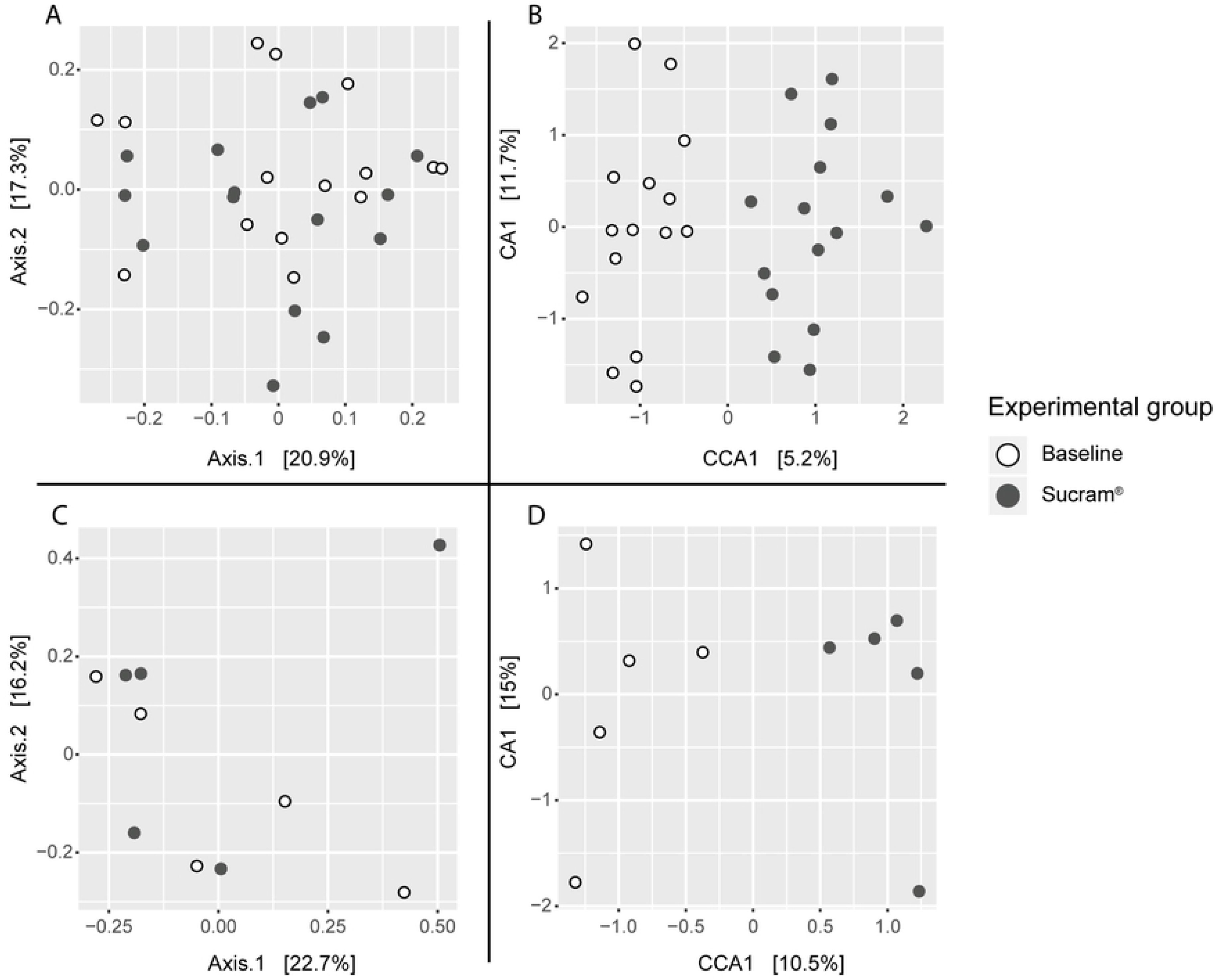
Beta diversity of rumen microbial communities in response to Sucram ^®^ addition to the diet. Principal coordinate analysis of rumen epithelium (A) and rumen content (C) samples. Canonical coordinate analysis of rumen epithelium (B) and rumen content (D) microbial communities in response to Sucram^®^ addition. Data are shown for baseline (before) and after 28 days of Sucram^®^ addition to the diet. All plots are based on Bray Curtis differences.

Significant differences in abundance of 11 OTUs were identified using LEfSe. However, out of the 100 most abundant rumen content OTUs, none were found to be significantly different in abundance between the experimental groups (Supplementary Table S5).

### Rumen epithelium microbial communities

Overall, 6,511 OTUs were generated after quality control and removal of OTUs representing less than 10 sequences from the 3.05 million high quality sequences obtained from 30 samples. The average number of sequences per sample was 101,742, with a standard deviation of 18,577 sequences. Most of the reads (97.7%) were bacterial, 2.3% were archaeal. From the 6,511 OTUs, 26 phyla were identified with *Firmicutes*, *Bacteroidetes*, and *Proteobacteria* being the most abundant: representing 32%, 28%, and 21% of all reads, respectively (Fig. 1). The most abundant OTU within the epithelial data set was identified as *Ruminobacter* RUG14687 with 100% sequence similarity using the RGC and *Succinivibrionaceae*_UCG-001 (88.98% sequence similarity, Silva v132), and accounted for 11.7 % of all reads from the epithelial data set. OTUs 2, 3, 4 and 5 were classified as *Mogibacterium*, *Butyrivibrio*, *Campylobacter* and *Prevotella*, accounting for 3.0%, 1.7%, 1.7% and 1.4% of all epithelial reads, respectively. A list of the 50 most abundant rumen wall OTUs can be found in Supplementary Table S6. On genus level, the most abundant genera of the rumen wall dataset include: *Succinivibrionaceae*_UCG-001 (12%), *Prevotella_1* (8.7%), *Butyrivibrio_2* (6.1%), *Rikenella* (4.2%), *Treponema_2* (4%), and *Mogibacterium* (3.7%) (Fig. 2).

When comparing BL and Sucram^®^ epithelial experimental groups, we observed no significant differences in diversity (Shannon, *P* = 0.64, species richness (Chao, *P* = 0.07) and community evenness (Simpson, *P* = 0.73) (Supplementary Table S7). Similar to the rumen content dataset, no significant differences (p-value: 0.2, R-value: 0.025) were detected when comparing entire bacterial communities of experimental groups using ANOSIM. This result was corroborated by the lack of apparent clustering of experimental groups seen in the PCoA (Fig. 3). BL and Sucram^®^ samples cluster separately in CCA (Fig. 3); however, only 5.2% of the total variation was due to experimental group.

When comparing individual OTUs using LEfSe, the abundances of 78 OTUs were significantly different between experimental groups. Among the 100 most abundant OTUs, 20 OTUs were significantly different in abundance abundant between experimental groups, with 10 OTUs found to be more abundant in the Sucram^®^ experimental group and 10 OTUs that showed higher abundance in BL samples (Fig. 4, Supplementary Table S8). The 10 OTUs found to be more abundant in the Sucram^®^ experimental group were classified as *Prevotellaceae* (OTUs 5, 11, 14, 33, 38, 80, 94), *Sharpea* (OTU 43) and *Bacteroidales* p-251-o5 (OTU 50, an uncultured member of the *Bacteroidetes*). The 10 OTUs more abundant in the BL samples were classified as *Methanomethylophilaceae* (OTU 31, 61, 71), *Methanobrevibacter* (OTU 21), *Desulfobulbus* (OTU 6, 34), *Rikenellaceae* RC9 gut group (OTU 26), *Treponema* (OTU 44), RBG-16-49-21 (OTU 47, a member of the *Leptospiraceae* family) and *Bacteroidales* F082 (OTU 53, an uncultured member of the *Bacteroidales*).

**Fig. 4.**
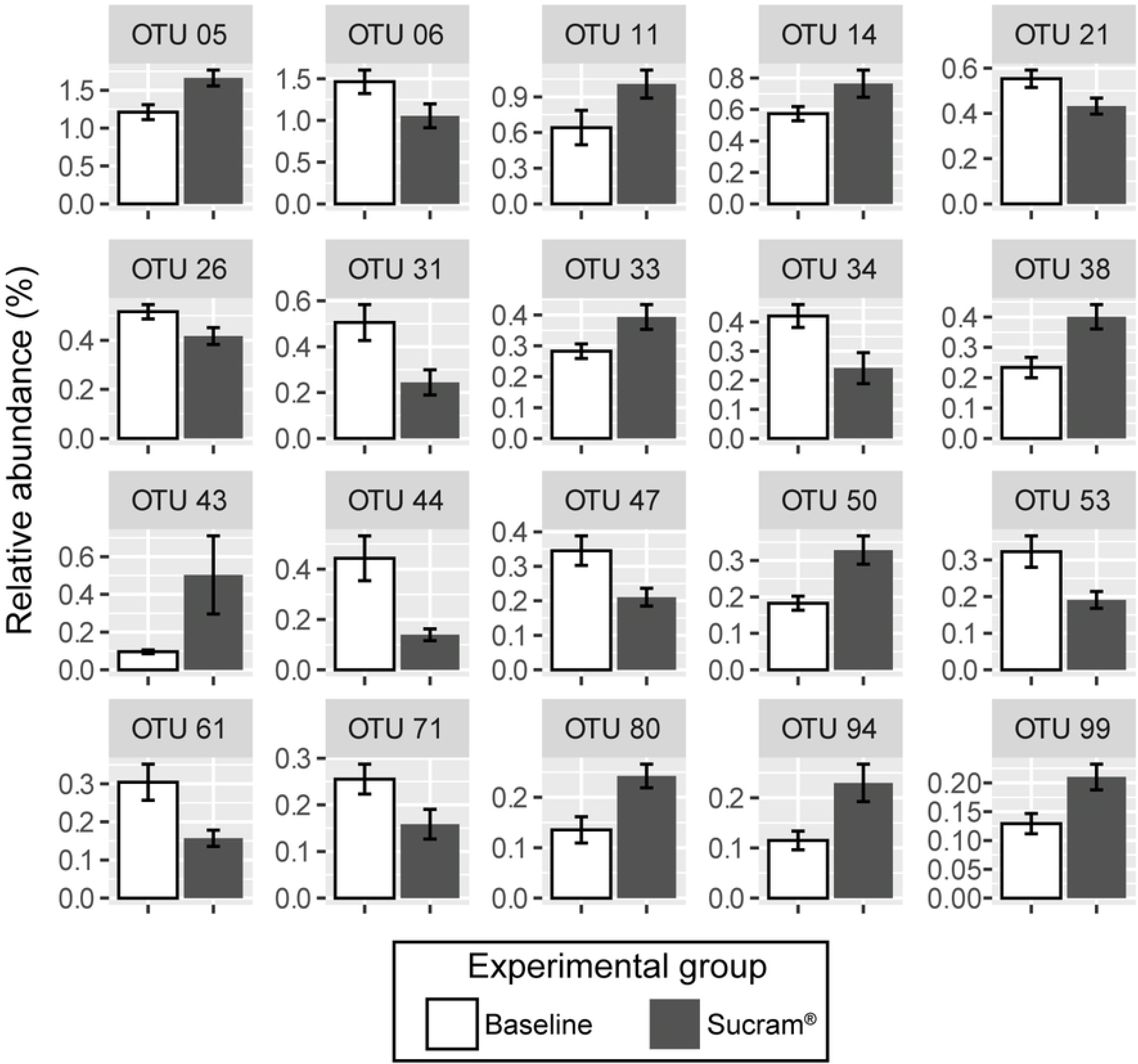
Relative abundance of significantly different OTUs between baseline and after 28 days of Sucram ^®^ supplementation within the rumen epithelium microbial communities. Significantly different OTUs were identified with LEfSe [33] and p-values lower than 0.05 were considered significant. Only significant OTUs within the 100 most abundant OTUs are shown. Error bars represent the standard error of the mean. See Supplementary Table S6 for more details.

### Predictions of functional potential

Based on the 97% sequence similarity threshold mentioned above, 32 of the 100 most abundant OTUs from the rumen content dataset had matches to sequences within the RGC. Similarly, 29 of the 100 most abundant OTUs from the rumen epithelium dataset had matches to sequences within the RGC. The associated taxonomy based on standard BLAST, the Silva reference database and the information provided in the RGC can be found in Supplementary Tables S3 and S6.

BLASTn comparisons of all the representative sequences for all significantly different OTUs (LEfSe) between experimental groups, including the significant, lowly abundant OTUs beyond the 100 most abundant, 4 of 11 significant OTUs from the rumen content dataset and 19 of 78 significant OTUs from the rumen epithelium dataset had highly similar (>97%) matches to the RGC. Results can be found in Supplementary Tables S5 and S8.

Several genes related to feed efficiency were identified in the annotated RGC genomes of the significant OTUs, however, no clear trend was identified in either the rumen content or rumen epithelium datasets (Supplementary Figures S1 and S2). Very few of the putative genes for saccharin degradation that were suggested by Deng et al. [39] were identified in any of the genomes analyzed, and those that were, were identified within both experimental groups. As mentioned above, the lack of knowledge pertaining to genes involved in saccharin degradation makes interpreting this data difficult, but does offer a unique opportunity for future research.

Within the rumen content set, genes involved in the lactate production pathway were identified in organisms belonging to both the Sucram and baseline experimental groups. Genes involved in propionate, valerate and isovalerate synthesis were only found in the Sucram experimental group only, however, there was only a single rumen content baseline OTU (717) that had a match within the RGC. A single gene involved in methanogenesis [E.C. 1.8.98.1] was found in the genome matching OTU 717 as well, which was more abundant in the baseline experimental group.

Within the rumen epithelium dataset, genes involved in the propionate and lactate production pathways were identified in organisms belonging to both the Sucram and baseline experimental groups. The aldehyde dehydrogenase (NAD^+^) [E.C. 1.2.1.3] within the valerate and isovalerate synthesis pathways was found in genomes related to organisms that were significantly more abundant in the Sucram experimental group. Butyrate kinase [E.C. 2.7.2.7], a key enzyme in the synthesis of butyrate, was only found in a single genome matching OTU260 that was significantly more abundant in the baseline experimental group. Genes involved in methanogenesis were identified in genomes related to OTU21 which was classified as a *Methanobrevibacter sp*. and was more abundant in the baseline experimental group.

## Discussion

Before discussing the results and implications of this study, we want to highlight some of the limitations. The main aim of our study was to identify if the presence of Sucram® has an influence or rumen microbial communities. To ensure the additive was indeed present within the rumen, Sucram® was administered directly through the fistula. Any sensory stimuli (i.e. taste or smell) would be limited, as the sweetener was not added to the feed. Thus, as mentioned above, we did not measure feed intake, average daily gain or milk yield in response to administration of Sucram® for this study.

Despite the limitations, this is the first study to provide insight into the effect of saccharin-based sweeteners such as Sucram^®^ on rumen microbial communities present within the content as well as on the rumen epithelium. Until now, the research conducted to identify the effects of artificial sweeteners on mammalian gastrointestinal tract microbial communities has been exclusively focused on monogastric animal models such a rats and pigs [4, 7, 40]. For example, a recent study provided evidence that certain bacteria degrade saccharin-based artificial sweeteners, indicating artificial sweeteners may not be non-caloric to the host if they are degraded to usable metabolic products [39]. Other studies focusing on the effect of Sucram^®^ supplementation have reported changes in microbial communities in monogastric livestock. Daly *et al.* reported that Sucram^®^ alters the abundance of several bacterial phylotypes in porcine intestinal microbial communities including *Lactobacillus*, *Ruminoccocaceae* and *Veillonellaceae* [41]. Additionally, Kelly et al. provided evidence that the addition of Sucram^®^ to the diet led to changes in the diversity of mucosal bacteria within the small intestine in swine, specifically a decrease in *Campylobacter coli* and an increase in members of the *Helicobacteraceae* family [3]. Additionally, a reduction of *Ruminococcus* was documented in a study analyzing the effect of saccharin on inflammatory molecules and gut dysbiosis in mice [40]. Although no significant differences were found for OTUs classified as *Lactobacillus, Veillonellaceae*, *Campylobacter* or *Helicobacteraceae,* the study reported in this manuscript observed a significant decrease in *Rumincoccaceae* (OTU 106) similar to other recent studies in monogastric animals [40, 42]. These studies and the study reported here demonstrate that Sucram^®^ C-150 does have an effect on mammalian gut microbial populations, but the changes in microbial community structure are largely distinct between monogastric and ruminant species.

In the present report, we show that the addition of Sucram^®^ did not induce significant changes in overall species richness, evenness or diversity (Supplementary Tables S3 and S5), in either rumen content or rumen epithelial microbial communities. This is somewhat unexpected, considering the impact Sucram^®^ supplementation had on highly abundant phylotypes, especially the highly abundant OTUs within the rumen epithelium data set (Supplementary Table S6). Although some members of the microbial communities have increased or decreased considerably in abundance (i.e. *Prevotellaceae*, *Desulfobulbaceae* and methanogenic archaea), there were no phylotypes that were completely diminished after the supplementation of Sucram^®^. This suggests that supplementation of Sucram^®^ does not have a major overall impact on the entire rumen microbial ecosystem, and is in contrast to observations reported in some monogastric animal studies which reported significant community differences in microbial communities due to the addition of saccharin-based sweeteners [4, 42]. It can be speculated that artificial sweeteners may specifically influence bacteria that can metabolize saccharin-based artificial sweeteners, which may also lead to secondary effects on rumen microbiota composition. As mentioned above, degradation of saccharin has been described for a number of bacteria, but so far, only for those obtained from environmental, non-animal samples [39]. This would also suggest that ruminant microbial communities might metabolize these sweeteners differently than monogastric microbial communities, although this requires verification in future studies.

Notably, the inclusion of Sucram^®^ affected the microbial populations of the rumen epithelium more strongly than those of the rumen content (Supplementary Table S6). Because it is well established that the microbial communities of the rumen content and epithelium are largely distinct, in both community composition and function, it is reasonable to assume that Sucram^®^ would not necessarily affect both communities to similar degrees and in comparable ways [17, 19, 20, 22, 43, 44]. However, the functional mechanisms behind why the rumen epithelial community appears more responsive to Sucram^®^ supplementation remain unknown. Given the importance of rumen epithelial microbial communities in key metabolic processes such as nutrient exchange (e.g. urea, sulfate reduction, and oxygen scavenging) between rumen content and the host animal [23, 45], and the importance of these interactions on the integrity of the rumen epithelium barrier function, this knowledge gap warrants future study.

We used the taxonomy generated based on NCBI BLAST, the Silva reference database and information provided in the RGC to compare our results with previously published work on related organisms. The consideration of differences between rumen epithelial experimental groups BL and Sucram^®^ with regards to significantly different OTUs may narrow the focus for future research, providing a list of potentially relevant organisms. Additionally, we can use previous research to infer possible function of these organisms, leading to potential metabolic markers for future research as well. However, functional prediction is not experimental proof of these pathways being expressed, so they should only be considered hypothesis. The comparisons of the OTUs and our interpretations of these results are listed below.

Two OTUs (6 and 34) with reduced abundance found in samples supplemented with Sucram^®^ were classified as potential sulfate-reducing bacteria (SRB) within the *Desulfobulbus* genus and were related to *Desulfobulbus oligotrophicus* (sequence similarity >97% using NCBI BLAST [46, 47]. SRB are commonly found within the gastrointestinal tract of ruminants [48] and particularly at the rumen wall [44, 49] and are known to reduce SO_3_^2−^ to SO_4_^2−^ and H_2_S [50]. This leads to the reduction of available metabolic hydrogen generated in cellulose and SCFA metabolism [47, 51]. Removal of metabolic hydrogen is thought to reduce the synthesis of methane, reducing the growth potential of methanogenic archaea [47, 50–52]. At the same time, H_2_S is considered cytotoxic and linked with inflammation within the intestine [53, 54]. It is therefore possible that a reduction of SRB could lead to increased methane production due to reduced competition for hydrogen. As a consequence, this could also result in increased levels of acetate and lactate, which could be used for milk production by the ruminant and reducing inflammation in the rumen and intestinal lining.

In addition to the decrease of these potential SRB, a decrease in methanogenic archaea was also observed. Supplementation of Sucram^®^ reduced the abundance of OTUs 21, 31, 61 and 71 which were all classified as potential methane-producing archaea (Figure 4). OTU 21 matched a genome from the RGC with 100% similarity and as expected, several of the genes involved in methanogenesis were identified. While this finding contradicts the hypothesis that SRBs directly compete with methanogens as mentioned above, it should be noted that the study by Abram et al [47] and the present study were conducted in two highly different ecosystems (waste water versus rumen). Because methane production is energetically costly and therefore reduces feed efficiency [52, 55–57], this decrease in methanogenic archaeal abundance suggests that a supplementation of Sucram^®^ may confer an overall benefit in feed efficiency to the host. Additionally, reduction in methanogenic archaea could also reduce the generation and release of methane into the atmosphere [52].

Sucram^®^ supplementation also decreased the abundance of OTU 26, which was classified as a member of the uncultivated *Rikenellaceae*_RC9_gut_group (Figure 4). There is little information available for members of this family and their possible function in ruminants, although they have been described in ruminants before [58]. It has been shown that some species that are closely related to the *Rikenellaceae* family can tolerate bile salts and produce succinate from glucose metabolism [59], but this may not be the case for the *Rikenellaceae* family specifically.

We also found that OTU 44, which was classified as *Treponema*, was significantly decreased in samples supplemented with Sucram^®^. In previous literature, *Treponema* species have often been associated with cellulolytic capabilities on the surface of plant material and are known to contain genes allowing the degradation of pectin, xylan, cellulose and starch [60, 61]. When grown in co-culture with other cellulose degraders (*Ruminococcus albus* and *Fibrobacter succinogenes*), *Treponema bryantii* produced lactic and succinic acids from the byproducts of cellulose degradation. However, the *Treponema* OTU found in this study (OTU 44) was <90% similar to *Treponema bryantii*.

The abundance of OTU 106, classified as *Ruminococcus*, was significantly decreased with the supplementation of Sucram^®^. Members of the genus *Ruminoccocus* are well recognized as cellulose degraders with the ability to produce butyrate from the fermentation of cellulose [62]. *Ruminoccocus* is linked to higher residual feed intake, suggesting lower feed efficiency in animals that harbor these organisms in high abundance [63]. Although such animals may be less feed efficient, producing butyrate is beneficial for the host as it positively stimulates the immune system and increases tight junction strength [64]. As a pilot study, this trial did not record feed intake or milk yield of the cows, so it is impossible to relate the reduction in *Ruminococcus* to feed efficiency and animal health. Additional research testing the effect of Sucram^®^ on *Ruminoccocus* species and how it relates to overall animal efficiency is warranted.

In contrast to the decrease of different phylotypes, Sucram^®^ supplementation led to an increase in abundance of several rumen epithelium OTUs classified as bacteria known to aid in digestibility. Several OTUs which were identified as potentially hemicellulolytic and proteolytic (*Prevotella,* OTUs 5, 11, 14, 33, and 38) were found to be in higher abundance in samples supplemented with Sucram^®^ in comparison to baseline samples. Additionally, some OTUs were related to bacteria known to produce SCFAs such as lactate, acetate, succinate and propionate (*Sharpea* and *Prevotella*, OTUs 3, 11, 14, 33, 38 and 43) [65–67]. This suggests that cattle exposed to Sucram^®^ may be more efficiently degrading non-cellulose substrates, thereby potentially increasing concentrations of certain SCFAs.

*Prevotella* is often considered the most abundant bacterial genus in ruminants [14, 68]. Indeed, *Prevotella* was the most abundant genus in both the rumen content and rumen wall datasets. *Prevotella* species are often associated with plant-rich diets, and they are known to produce a variety of SCFAs such as acetate, succinate, propionate, and valerate which are absorbed and utilized by the host animal [69].Three OTUs (5, 33 and 161) identified as *Prevotella* with differing abundances between experimental groups were matched with the RGC, and genes related to propionate (OTUs 5 and 33) and lactate (OTU 161) production were identified within the matching genomes. Furthermore, cattle with greater abundance of *Prevotella* had lower residual feed intake and higher feed efficiency [63]. Finally, the abundance of *Prevotella* has been demonstrated to increase in goats fed higher levels of concentrate, a finding likely attributable to *Prevotella* species’ ability to utilize starch for energy production [70]. In the present study, it may be the case that *Prevotella* was capable of harnessing Sucram^®^ as an energy source, thereby promoting its growth. This warrants future research investigating the effect of Sucram^®^ on *Prevotella* species in regards to feed efficiency.

In this study, many highly abundant OTUs classified within the genus *Prevotella*, as well as OTU 43 (classified as *Sharpea azabuensis)* were found to be in higher abundance in Sucram^®^-supplemented samples (Figure 4). As previously stated, both *Prevotella* and *Sharpea* are known SCFA producers. A recent amplicon sequencing study established a link between lactate-producing bacteria, *Sharpea azabuensis*, and succinate and propionate-producing bacteria *Prevotella bryantii* where it was discovered sheep with lower methane emissions had a higher abundance of *Sharpea* and *Prevotella*. [56]. A follow-up study compared the metagenomes and metatranscriptomes of rumen microbiota from sheep with either high or low methane yields, further demonstrating that *Sharpea* reduces available metabolic hydrogen during SCFA synthesis thereby reducing methane production, resulting in a higher feed efficiency [57]. This can be connected to saccharin-based sweeteners such as Sucram^®^ through work by Suez et al, who identified increased expression of genes in glycogen degradation leading to increased production of propionate and acetate in rodents fed saccharin sweeteners [4]. Additionally, the inclusion of Sucram^®^ was shown to increase levels of lactate in the gastrointestinal lumen of pigs [41]. Bacteria producing SCFAs are in direct competition with methanogenic archaea [51, 52, 71], which could explain the reduction in methanogen abundance observed in this study. It can be hypothesized that Sucram^®^ might be able to modify the composition of the rumen microbiota in a way that decreases methane production by promoting growth SCFA producing bacteria, leading to an increase in livestock feed efficiency and productivity.

Additionally, to identify genes that may have implications on feed efficiency which Sucram^®^ differentially influences, we mined known rumen genomes that matched some of the 16S rRNA genes from this study with more than 97% sequence similarity for genes that may have implications on both feed efficiency and saccharin degradation. 36% and 24% of all significant OTUs for the rumen content and rumen epithelium respectively, had 16S rRNA gene sequences matches to the RGC. The results of this analytical method did offer additional supportive information for statements made above, such as confirming the presence of genes involved in lactate and propionate production in *Sharpea azabuensis* and *Prevotella* sp. genomes, genes involved in methane production in *Methanobrevibacter* sp. genomes, etc. It also provided potentially new implications for future research, such as implicating *Sharpea azabuensis* possible involvement in valerate and isovalerate production. Finally, it may offer functional predictions for previously uncultured organisms important to this study by matching 16S rRNA genes with rumen reference genomes assembled in a number of recent studies [34, 35].

Comparisons of OTUs using the RGC as a reference are a prediction of genetic potential. Using these predictions to suggest function should only be viewed as a hypothesis until experimentally tested. However, as seen in [72, 73], genomic prediction based on 16S rRNA gene sequence similarity can be useful in identifying gene or metabolic targets in future studies. And, as the research into rumen bacterial communities and their function progresses, the usefulness of such habitat-specific reference databases such as the RGC should increase. Identifying and including genes important in cellulose degradation, SCFA synthesis, vitamin synthesis, nitrogen cycling, and other metabolically important processes for ruminants would improve predictions of genomic potential and detection of pathways relevant to the experimental treatment. Genes involved in saccharin degradation and utilization are wholly absent from the literature as well, which could have implications on animal health and performance.

## Conclusion

This pilot study demonstrated that the rumen microbial community remains largely stable in response to the inclusion of Sucram^®^, with only a few significant changes in abundance of certain microbial phylotypes. A prediction of the important phylotypes’ genetic potential identified the presence of key genes related to feed efficiency. The differences observed in response to Sucram^®^ supplementation were most profound in the rumen epithelial microbial communities, which could have important implications for the integrity of the rumen epithelium to serve as a barrier preventing inflammation and infiltration of pathogens. OTUs decreased after Sucram^®^ supplementation were linked to methane production and reduced feed efficiency, whereas OTUs increased following Sucram^®^ supplementation were associated with propionate and lactate production potential and increased feed efficiency. Therefore, supplementation with Sucram^®^ may foster a microbial community that decreases available metabolic hydrogen for methane generation and produces SCFAs important for ruminant performance. This in turn may lead to more efficient ruminant livestock with lower methane emission. However, we also want to highlight the preliminary nature of the current pilot study and the need for future validations of our findings. In conclusion, this pilot study suggests that the supplementation of saccharin-based sweeteners such as Sucram^®^ causes novel and potentially beneficial effects on ruminant health and performance other than increased palatability, possibly driven by changes in the rumen microbiota especially the microbial communities of the lesser-studied rumen epithelium.

## Acknowledgements

We would like to acknowledge Dr. Alexandra Blanchard, Pancosma S.A./ADM Group, Rolle, Switzerland, for critically reading the manuscript.

## Supporting information captions

**Supplementary Table S1:** Dietary composition

**Supplementary Table S2:** Genes related to feed efficiency and saccharin degradation. Genes related to feed efficiency were based on Shabat et al. 2016, genes suggested to be involved in saccharin degradation are based on Deng et al. 2019.

**Supplementary Table S3:** The 50 most abundant OTUs within rumen content samples

**Supplementary Table S4:** Chao species richness, Shannon and non-parametric Shannon Diversity, and Simpson evenness comparisons between Sucram and baseline experimental groups within the rumen content samples.

**Supplementary Table S5:** Significantly different rumen content OTUs between Sucram and baseline determined with LEfSe.

**Supplementary Table S6:** The 50 most abundant OTUs within rumen epithelial samples

**Supplementary Table S7:** Chao species richness, Shannon and non-parametric Shannon Diversity, and Simpson evenness comparisons between Sucram and baseline experimental groups within the rumen epithelium samples.

**Supplementary Table S8:** Significantly different rumen epithelium OTUs between Sucram and baseline determined with LEfSe.

**Supplementary Figure S1.** Predicted genetic potential of rumen content organisms that significantly differ in abundance between experimental groups, using matching published genomes.

**Supplementary Figure S2.** Predicted genetic potential of rumen epithelium organisms that significantly differ in abundance between experimental groups, using matching published genomes.

## References

1. McMeniman JP, Rivera JD, Schlegel P, Rounds W, Galyean ML. Effects of an artificial sweetener on health, performance, and dietary preference of feedlot cattle. J Anim Sci. 2006;84(9):2491–500. Epub 2006/08/16. doi: 10.2527/jas.2006-098. PubMed PMID: 16908654.

2. Ponce CH, Brown MS, Silva JS, Schlegel P, Rounds W, Hallford DM. Effects of a dietary sweetener on growth performance and health of stressed beef calves and on diet digestibility and plasma and urinary metabolite concentrations of healthy calves. J Anim Sci. 2014;92(4):1630–8. Epub 2014/03/26. doi: 10.2527/jas.2013-6795. PubMed PMID: 24663208.

3. Kelly J, Daly K, Moran AW, Ryan S, Bravo D, Shirazi-Beechey SP. Composition and diversity of mucosa-associated microbiota along the entire length of the pig gastrointestinal tract; dietary influences. Environ Microbiol. 2017;19(4):1425–38. Epub 2016/11/22. doi: 10.1111/1462-2920.13619. PubMed PMID: 27871148.

4. Suez J, Korem T, Zeevi D, Zilberman-Schapira G, Thaiss CA, Maza O, et al. Artificial sweeteners induce glucose intolerance by altering the gut microbiota. Nature. 2014;514(7521):181–6. Epub 2014/09/19. doi: 10.1038/nature13793. PubMed PMID: 25231862.

5. Suez J, Korem T, Zilberman-Schapira G, Segal E, Elinav E. Non-caloric artificial sweeteners and the microbiome: findings and challenges. Gut Microbes. 2015;6(2):149–55. Epub 2015/04/02. doi: 10.1080/19490976.2015.1017700. PubMed PMID: 25831243; PubMed Central PMCID: PMCPMC4615743.

6. Abou-Donia MB, El-Masry EM, Abdel-Rahman AA, McLendon RE, Schiffman SS. Splenda alters gut microflora and increases intestinal p-glycoprotein and cytochrome p-450 in male rats. J Toxicol Environ Health A. 2008;71(21):1415–29. Epub 2008/09/19. doi: 10.1080/15287390802328630. PubMed PMID: 18800291.

7. Anderson RL, Kirkland JJ. The effect of sodium saccharin in the diet on caecal microflora. Food Cosmet Toxicol. 1980;18(4):353–5. Epub 1980/08/01. PubMed PMID: 7007181.

8. Frankenfeld CL, Sikaroodi M, Lamb E, Shoemaker S, Gillevet PM. High-intensity sweetener consumption and gut microbiome content and predicted gene function in a cross-sectional study of adults in the United States. Ann Epidemiol. 2015;25(10):736–42 e4. Epub 2015/08/15. doi: 10.1016/j.annepidem.2015.06.083. PubMed PMID: 26272781.

9. Ruiz-Ojeda FJ, Plaza-Diaz J, Saez-Lara MJ, Gil A. Effects of Sweeteners on the Gut Microbiota: A Review of Experimental Studies and Clinical Trials. Adv Nutr. 2019;10(suppl_1):S31–S48. Epub 2019/02/06. doi: 10.1093/advances/nmy037. PubMed PMID: 30721958; PubMed Central PMCID: PMCPMC6363527.

10. Bokulich NA, Blaser MJ. A bitter aftertaste: unintended effects of artificial sweeteners on the gut microbiome. Cell Metab. 2014;20(5):701–3. Epub 2014/12/03. doi: 10.1016/j.cmet.2014.10.012. PubMed PMID: 25440050; PubMed Central PMCID: PMCPMC4494042.

11. Labrecque MT, Malone D, Caldwell KE, Allan AM. Impact of Ethanol and Saccharin on Fecal Microbiome in Pregnant and Non-Pregnant Mice. J Pregnancy Child Health. 2015;2(5). Epub 2016/03/19. doi: 10.4172/2376-127X.1000193. PubMed PMID: 26989786; PubMed Central PMCID: PMCPMC4792281.

12. Paz HA, Hales KE, Wells JE, Kuehn LA, Freetly HC, Berry ED, et al. Rumen bacterial community structure impacts feed efficiency in beef cattle. J Anim Sci. 2018;96(3):1045–58. Epub 2018/04/05. doi: 10.1093/jas/skx081. PubMed PMID: 29617864; PubMed Central PMCID: PMCPMC6093515.

13. Krause DO, Denman SE, Mackie RI, Morrison M, Rae AL, Attwood GT, et al. Opportunities to improve fiber degradation in the rumen: microbiology, ecology, and genomics. FEMS Microbiol Rev. 2003;27(5):663–93. Epub 2003/11/26. doi: 10.1016/S0168-6445(03)00072-X. PubMed PMID: 14638418.

14. Bekele AZ, Koike S, Kobayashi Y. Genetic diversity and diet specificity of ruminal Prevotella revealed by 16S rRNA gene-based analysis. FEMS Microbiology Letters. 2010;305(1):49–57. doi: 10.1111/j.1574-6968.2010.01911.x.

15. Plaizier JC, Krause DO, Gozho GN, McBride BW. Subacute ruminal acidosis in dairy cows: The physiological causes, incidence and consequences. VET J. 2008;176(1):21–31. doi: https://doi.org/10.1016/j.tvjl.2007.12.016.

16. Steele MA, Croom J, Kahler M, AlZahal O, Hook SE, Plaizier K, et al. Bovine rumen epithelium undergoes rapid structural adaptations during grain-induced subacute ruminal acidosis. Am J Physiol Regul Integr Comp Physiol. 2011;300(6):R1515–R23. doi: 10.1152/ajpregu.00120.2010. PubMed PMID: 21451145.

17. Liu JH, Zhang ML, Zhang RY, Zhu WY, Mao SY. Comparative studies of the composition of bacterial microbiota associated with the ruminal content, ruminal epithelium and in the faeces of lactating dairy cows. Microb Biotechnol. 2016;9(2):257–68. Epub 2016/02/03. doi: 10.1111/1751-7915.12345. PubMed PMID: 26833450; PubMed Central PMCID: PMCPMC4767291.

18. Tamate H, Kikuchi T, Onodera A, Nagatani T. Scanning electron microscopic observation on the surface structure of the bovine rumen mucosa. Arch Histol Jpn. 1971;33(4):273–82. Epub 1971/10/01. PubMed PMID: 5168659.

19. McCowan RP, Cheng KJ, Bailey CB, Costerton JW. Adhesion of bacteria to epithelial cell surfaces within the reticulo-rumen of cattle. Appl Environ Microbiol. 1978;35(1):149–55. Epub 1978/01/01. PubMed PMID: 623459; PubMed Central PMCID: PMCPMC242795.

20. Cheng KJ, McCowan RP, Costerton JW. Adherent epithelial bacteria in ruminants and their roles in digestive tract function. Am J Clin Nutr. 1979;32(1):139–48. Epub 1979/01/01. doi: 10.1093/ajcn/32.1.139. PubMed PMID: 367141.

21. Chen Y, Penner GB, Li M, Oba M, Guan LL. Changes in bacterial diversity associated with epithelial tissue in the beef cow rumen during the transition to a high-grain diet. Appl Environ Microbiol. 2011;77(16):5770–81. Epub 2011/06/28. doi: 10.1128/AEM.00375-11. PubMed PMID: 21705529; PubMed Central PMCID: PMCPMC3165274.

22. Chen Y, Oba M, Guan LL. Variation of bacterial communities and expression of Toll-like receptor genes in the rumen of steers differing in susceptibility to subacute ruminal acidosis. Vet Microbiol. 2012;159(3-4):451–9. Epub 2012/05/25. doi: 10.1016/j.vetmic.2012.04.032. PubMed PMID: 22622335.

23. Mann E, Wetzels SU, Wagner M, Zebeli Q, Schmitz-Esser S. Metatranscriptome Sequencing Reveals Insights into the Gene Expression and Functional Potential of Rumen Wall Bacteria. Front Microbiol. 2018;9:43–. doi: 10.3389/fmicb.2018.00043. PubMed PMID: 29410661.

24. Bryant MP. Bacterial species of the rumen. Bacteriol Rev. 1959;23(3):125–53. Epub 1959/09/01. PubMed PMID: 13805451; PubMed Central PMCID: PMCPMC181027.

25. White BA, Lamed R, Bayer EA, Flint HJ. Biomass Utilization by Gut Microbiomes. Annu. Rev. Microbiol. 2014;68(1):279–96. doi: 10.1146/annurev-micro-092412-155618. PubMed PMID: 25002092.

26. Parada AE, Needham DM, Fuhrman JA. Every base matters: assessing small subunit rRNA primers for marine microbiomes with mock communities, time series and global field samples. Environ Microbiol. 2016;18(5):1403–14. Epub 2015/08/15. doi: 10.1111/1462-2920.13023. PubMed PMID: 26271760.

27. Apprill A, McNally S, Parsons R, Weber L. Minor revision to V4 region SSU rRNA 806R gene primer greatly increases detection of SAR11 bacterioplankton. AQUAT MICROB ECOL. 2015;75. doi: 10.3354/ame01753.

28. Kozich JJ, Westcott SL, Baxter NT, Highlander SK, Schloss PD. Development of a dual-index sequencing strategy and curation pipeline for analyzing amplicon sequence data on the MiSeq Illumina sequencing platform. Appl Environ Microbiol. 2013;79(17):5112–20. Epub 2013/06/25. doi: 10.1128/AEM.01043-13. PubMed PMID: 23793624; PubMed Central PMCID: PMCPMC3753973.

29. R Core Team. R: A Language and Environment for Statistical Computing. Vienna, Austria2019.

30. Wickham H. ggplot2: Elegant Graphics for Data Analysis. Springer-Verlag New York; 2016.

31. McMurdie PJ, Holmes S. phyloseq: an R package for reproducible interactive analysis and graphics of microbiome census data. PLoS One. 2013;8(4):e61217. Epub 2013/05/01. doi: 10.1371/journal.pone.0061217. PubMed PMID: 23630581; PubMed Central PMCID: PMCPMC3632530.

32. Oksanen J, Blanchet FG, Friendly M, Kindt R, Legendre P, McGlinn D, et al. vegan: Community Ecology Package. 2019.

33. Segata N, Izard J, Waldron L, Gevers D, Miropolsky L, Garrett WS, et al. Metagenomic biomarker discovery and explanation. Genome Biol. 2011;12(6):R60. Epub 2011/06/28. doi: 10.1186/gb-2011-12-6-r60. PubMed PMID: 21702898; PubMed Central PMCID: PMCPMC3218848.

34. Stewart RD, Auffret MD, Warr A, Walker AW, Roehe R, Watson M. Compendium of 4,941 rumen metagenome-assembled genomes for rumen microbiome biology and enzyme discovery. Nat Biotechnol. 2019;37(8):953–61. Epub 2019/08/04. doi: 10.1038/s41587-019-0202-3. PubMed PMID: 31375809; PubMed Central PMCID: PMCPMC6785717.

35. Seshadri R, Leahy SC, Attwood GT, Teh KH, Lambie SC, Cookson AL, et al. Cultivation and sequencing of rumen microbiome members from the Hungate1000 Collection. Nat Biotechnol. 2018;36(4):359–67. Epub 2018/03/20. doi: 10.1038/nbt.4110. PubMed PMID: 29553575; PubMed Central PMCID: PMCPMC6118326.

36. Camacho C, Coulouris G, Avagyan V, Ma N, Papadopoulos J, Bealer K, et al. BLAST+: architecture and applications. BMC Bioinformatics. 2009;10:421. Epub 2009/12/17. doi: 10.1186/1471-2105-10-421. PubMed PMID: 20003500; PubMed Central PMCID: PMCPMC2803857.

37. Wattam AR, Davis JJ, Assaf R, Boisvert S, Brettin T, Bun C, et al. Improvements to PATRIC, the all-bacterial Bioinformatics Database and Analysis Resource Center. Nucleic Acids Res. 2017;45(D1):D535–D42. Epub 2016/12/03. doi: 10.1093/nar/gkw1017. PubMed PMID: 27899627; PubMed Central PMCID: PMCPMC5210524.

38. Shabat SK, Sasson G, Doron-Faigenboim A, Durman T, Yaacoby S, Berg Miller ME, et al. Specific microbiome-dependent mechanisms underlie the energy harvest efficiency of ruminants. ISME J. 2016;10(12):2958–72. Epub 2016/05/07. doi: 10.1038/ismej.2016.62. PubMed PMID: 27152936; PubMed Central PMCID: PMCPMC5148187.

39. Deng Y, Wang Y, Xia Y, Zhang AN, Zhao Y, Zhang T. Genomic resolution of bacterial populations in saccharin and cyclamate degradation. Sci Total Environ. 2019;658:357–66. Epub 2018/12/24. doi: 10.1016/j.scitotenv.2018.12.162. PubMed PMID: 30579193.

40. Bian X, Tu P, Chi L, Gao B, Ru H, Lu K. Saccharin induced liver inflammation in mice by altering the gut microbiota and its metabolic functions. Food Chem Toxicol. 2017;107(Pt B):530–9. Epub 2017/05/05. doi: 10.1016/j.fct.2017.04.045. PubMed PMID: 28472674; PubMed Central PMCID: PMCPMC5647777.

41. Daly K, Darby AC, Hall N, Wilkinson MC, Pongchaikul P, Bravo D, et al. Bacterial sensing underlies artificial sweetener-induced growth of gut *Lactobacillus*. Environ Microbiol. 2016;18(7):2159–71. Epub 2015/06/11. doi: 10.1111/1462-2920.12942. PubMed PMID: 26058469.

42. Daly K, Darby AC, Hall N, Nau A, Bravo D, Shirazi-Beechey SP. Dietary supplementation with lactose or artificial sweetener enhances swine gut *Lactobacillus* population abundance. Br J Nutr. 2014;111 Suppl 1:S30–5. Epub 2014/01/03. doi: 10.1017/S0007114513002274. PubMed PMID: 24382146.

43. Mead LJ, Jones GA. Isolation and presumptive identification of adherent epithelial bacteria (“epimural” bacteria) from the ovine rumen wall. Appl Environ Microbiol. 1981;41(4):1020–8. Epub 1981/04/01. PubMed PMID: 7195191; PubMed Central PMCID: PMCPMC243851.

44. Mao S, Zhang M, Liu J, Zhu W. Characterising the bacterial microbiota across the gastrointestinal tracts of dairy cattle: membership and potential function. Scientific reports. 2015;5:16116–. doi: 10.1038/srep16116. PubMed PMID: 26527325.

45. Jin D, Zhao S, Zheng N, Bu D, Beckers Y, Denman SE, et al. Differences in Ureolytic Bacterial Composition between the Rumen Digesta and Rumen Wall Based on ureC Gene Classification. Front Microbiol. 2017;8:385–. doi: 10.3389/fmicb.2017.00385. PubMed PMID: 28326079.

46. El Houari A, Ranchou-Peyruse M, Ranchou-Peyruse A, Dakdaki A, Guignard M, Idouhammou L, et al. *Desulfobulbus oligotrophicus sp. nov*., a sulfate-reducing and propionate-oxidizing bacterium isolated from a municipal anaerobic sewage sludge digester. Int J Syst Evol Microbiol. 2017;67(2):275–81. Epub 2016/12/03. doi: 10.1099/ijsem.0.001615. PubMed PMID: 27902225.

47. Abram JW, Nedwell DB. Inhibition of methanogenesis by sulphate reducing bacteria competing for transferred hydrogen. Arch Microbiol. 1978;117(1):89–92. Epub 1978/04/27. PubMed PMID: 678014.

48. Coleman GS. A sulphate-reducing bacterium from the sheep rumen. J Gen Microbiol. 1960;22:423–36. Epub 1960/04/01. doi: 10.1099/00221287-22-2-423. PubMed PMID: 13811147.

49. Wetzels SU, Mann E, Metzler-Zebeli BU, Pourazad P, Qumar M, Klevenhusen F, et al. Epimural Indicator Phylotypes of Transiently-Induced Subacute Ruminal Acidosis in Dairy Cattle. Front Microbiol. 2016;7:274. Epub 2016/03/15. doi: 10.3389/fmicb.2016.00274. PubMed PMID: 26973642; PubMed Central PMCID: PMCPMC4777738.

50. Isa Z, Grusenmeyer S, Verstraete W. Sulfate reduction relative to methane production in high-rate anaerobic digestion: microbiological aspects. Appl Environ Microbiol. 1986;51(3):580–7. Epub 1986/03/01. PubMed PMID: 16347019; PubMed Central PMCID: PMCPMC238922.

51. Jeyanathan J, Martin C, Morgavi DP. The use of direct-fed microbials for mitigation of ruminant methane emissions: a review. Animal. 2014;8(2):250–61. Epub 2013/11/28. doi: 10.1017/S1751731113002085. PubMed PMID: 24274095.

52. McAllister TA, Newbold CJ. Redirecting rumen fermentation to reduce methanogenesis. Australian Journal of Experimental Agriculture. 2008;48(2):7–13. doi: https://doi.org/10.1071/EA07218.

53. Nava GM, Carbonero F, Ou J, Benefiel AC, O'Keefe SJ, Gaskins HR. Hydrogenotrophic microbiota distinguish native Africans from African and European Americans. Environ Microbiol Rep. 2012;4(3):307–15. Epub 2013/06/14. doi: 10.1111/j.1758-2229.2012.00334.x. PubMed PMID: 23760794; PubMed Central PMCID: PMCPMC4258901.

54. Loubinoux J, Bronowicki JP, Pereira IA, Mougenel JL, Faou AE. Sulfate-reducing bacteria in human feces and their association with inflammatory bowel diseases. FEMS Microbiol Ecol. 2002;40(2):107–12. Epub 2002/05/01. doi: 10.1111/j.1574-6941.2002.tb00942.x. PubMed PMID: 19709217.

55. Tapio I, Snelling TJ, Strozzi F, Wallace RJ. The ruminal microbiome associated with methane emissions from ruminant livestock. J Anim Sci Biotechnol. 2017;8:7. Epub 2017/01/27. doi: 10.1186/s40104-017-0141-0. PubMed PMID: 28123698; PubMed Central PMCID: PMCPMC5244708.

56. Kittelmann S, Pinares-Patino CS, Seedorf H, Kirk MR, Ganesh S, McEwan JC, et al. Two different bacterial community types are linked with the low-methane emission trait in sheep. PLoS One. 2014;9(7):e103171. Epub 2014/08/01. doi: 10.1371/journal.pone.0103171. PubMed PMID: 25078564; PubMed Central PMCID: PMCPMC4117531.

57. Kamke J, Kittelmann S, Soni P, Li Y, Tavendale M, Ganesh S, et al. Rumen metagenome and metatranscriptome analyses of low methane yield sheep reveals a *Sharpea*-enriched microbiome characterised by lactic acid formation and utilisation. Microbiome. 2016;4(1):56. Epub 2016/10/21. doi: 10.1186/s40168-016-0201-2. PubMed PMID: 27760570; PubMed Central PMCID: PMCPMC5069950.

58. Schären M, Drong C, Kiri K, Riede S, Gardener M, Meyer U, et al. Differential effects of monensin and a blend of essential oils on rumen microbiota composition of transition dairy cows. Journal of Dairy Science. 2017;100(4):2765–83. doi: https://doi.org/10.3168/jds.2016-11994.

59. Abe K, Ueki A, Ohtaki Y, Kaku N, Watanabe K, Ueki K. *Anaerocella delicata gen. nov., sp. nov*., a strictly anaerobic bacterium in the phylum *Bacteroidetes* isolated from a methanogenic reactor of cattle farms. J Gen Appl Microbiol. 2012;58(6):405–12. doi: 10.2323/jgam.58.405.

60. Kudo H, Cheng KJ, Costerton JW. Interactions between Treponema bryantii and cellulolytic bacteria in the in vitro degradation of straw cellulose. Can J Microbiol. 1987;33(3):244–8. doi: 10.1139/m87-041.

61. Svartstrom O, Alneberg J, Terrapon N, Lombard V, de Bruijn I, Malmsten J, et al. Ninety-nine de novo assembled genomes from the moose (Alces alces) rumen microbiome provide new insights into microbial plant biomass degradation. ISME J. 2017;11(11):2538–51. Epub 2017/07/22. doi: 10.1038/ismej.2017.108. PubMed PMID: 28731473; PubMed Central PMCID: PMCPMC5648042.

62. Takahashi K, Nishida A, Fujimoto T, Fujii M, Shioya M, Imaeda H, et al. Reduced Abundance of Butyrate-Producing Bacteria Species in the Fecal Microbial Community in Crohn's Disease. Digestion. 2016;93(1):59–65. doi: 10.1159/000441768.

63. Li F, Guan LL. Metatranscriptomic Profiling Reveals Linkages between the Active Rumen Microbiome and Feed Efficiency in Beef Cattle. Appl Environ Microbiol. 2017;83(9). Epub 2017/02/27. doi: 10.1128/AEM.00061-17. PubMed PMID: 28235871; PubMed Central PMCID: PMCPMC5394315.

64. Peng L, Li Z-R, Green RS, Holzman IR, Lin J. Butyrate enhances the intestinal barrier by facilitating tight junction assembly via activation of AMP-activated protein kinase in Caco-2 cell monolayers. J Nutr. 2009;139(9):1619–25. Epub 2009/07/22. doi: 10.3945/jn.109.104638. PubMed PMID: 19625695.

65. Kumar S, Treloar BP, Teh KH, McKenzie CM, Henderson G, Attwood GT, et al. Sharpea and Kandleria are lactic acid producing rumen bacteria that do not change their fermentation products when co-cultured with a methanogen. Anaerobe. 2018;54:31–8. Epub 2018/07/29. doi: 10.1016/j.anaerobe.2018.07.008. PubMed PMID: 30055268.

66. Pybus V, Onderdonk AB. The Effect of pH on Growth and Succinate Production by Prevotella bivia. Microbial Ecology in Health and Disease. 1996;9(1):19–25. doi: 10.3109/08910609609167725.

67. Strobel HJ. Vitamin B12-dependent propionate production by the ruminal bacterium Prevotella ruminicola 23. Appl Environ Microbiol. 1992;58(7):2331–3. Epub 1992/07/01. PubMed PMID: 1637169; PubMed Central PMCID: PMCPMC195777.

68. Henderson G, Cox F, Ganesh S, Jonker A, Young W, Global Rumen Census C, et al. Rumen microbial community composition varies with diet and host, but a core microbiome is found across a wide geographical range. Sci Rep. 2015;5:14567. Epub 2015/10/10. doi: 10.1038/srep14567. PubMed PMID: 26449758; PubMed Central PMCID: PMCPMC4598811.

69. Emerson EL, Weimer PJ. Fermentation of model hemicelluloses by Prevotella strains and Butyrivibrio fibrisolvens in pure culture and in ruminal enrichment cultures. Appl Microbiol Biotechnol. 2017;101(10):4269–78. Epub 2017/02/10. doi: 10.1007/s00253-017-8150-7. PubMed PMID: 28180916.

70. Han X, Li B, Wang X, Chen Y, Yang Y. Effect of dietary concentrate to forage ratios on ruminal bacterial and anaerobic fungal populations of cashmere goats. Anaerobe. 2019;59:118–25. Epub 2019/06/23. doi: 10.1016/j.anaerobe.2019.06.010. PubMed PMID: 31228671.

71. Hill J, McSweeney C, Wright AG, Bishop-Hurley G, Kalantar-Zadeh K. Measuring Methane Production from Ruminants. Trends Biotechnol. 2016;34(1):26–35. Epub 2015/11/26. doi: 10.1016/j.tibtech.2015.10.004. PubMed PMID: 26603286.

72. Langille MG, Zaneveld J, Caporaso JG, McDonald D, Knights D, Reyes JA, et al. Predictive functional profiling of microbial communities using 16S rRNA marker gene sequences. Nat Biotechnol. 2013;31(9):814–21. Epub 2013/08/27. doi: 10.1038/nbt.2676. PubMed PMID: 23975157; PubMed Central PMCID: PMCPMC3819121.

73. Wilkinson TJ, Huws SA, Edwards JE, Kingston-Smith AH, Siu-Ting K, Hughes M, et al. CowPI: A Rumen Microbiome Focussed Version of the PICRUSt Functional Inference Software. Front Microbiol. 2018;9:1095. Epub 2018/06/12. doi: 10.3389/fmicb.2018.01095. PubMed PMID: 29887853; PubMed Central PMCID: PMCPMC5981159.

